# When to Trust Epigenetic Clocks: Avoiding False Positives in Aging Interventions

**DOI:** 10.1101/2024.10.22.619720

**Authors:** Daniel S. Borrus, Raghav Sehgal, Jenel Fraij Armstrong, Jessica Kasamoto, John Gonzalez, Albert Higgins-Chen

**Affiliations:** Department of Psychiatry, Yale University School of Medicine, New Haven, CT, USA; Program in Computational Biology and Bioinformatics, Yale University School of Medicine, New Haven, CT, USA; Department of Pathology, Yale University School of Medicine, New Haven, CT, USA

**Keywords:** epigenetic clocks, biomarkers, aging interventions, age reversal, false positives

## Abstract

Recent human studies have suggested that aging interventions can reduce aging biomarkers related to morbidity and mortality risk. Such biomarkers may potentially serve as early, rapid indicators of effects on healthspan. An increasing number of studies are measuring intervention effects on epigenetic clocks, commonly used aging biomarkers based on DNA methylation profiles. However, with dozens of clocks to choose from, different clocks may not agree on the effect of an intervention. Furthermore, changes in some clocks may simply be the result of technical noise causing a false positive result. To address these issues, we measured the variability between 6 popular epigenetic clocks across a range of longitudinal datasets containing either an aging intervention or an age-accelerating event. We further compared them to the same clocks re-trained to have high test-retest reliability. We find the newer generation of clocks, trained on mortality or rate-of-aging, capture aging events more reliably than those clocks trained on chronological age, as these show consistent effects (or lack thereof) across multiple clocks including high-reliability versions, and including after multiple testing correction. In contrast, clocks trained on chronological age frequently show sporadic changes that are not replicable when using high-reliability versions of those same clocks, or when using newer generations of clocks and these results do not survive multiple-testing correction. These are likely false positive results, and we note that some of these clock changes were previously published, suggesting the literature should be re-examined. This work lays the foundation for future clinical trials that aim to measure aging interventions with epigenetic clocks, by establishing when to attribute a given change in biological age to a *bona fide* change in the aging process.

## Introduction

In the pursuit of extending human healthspan, various interventions — such as dietary regimens, supplements, and pharmaceutical agents — are being developed to target the underlying biological mechanisms associated with aging (López-Otín et al. 2023; Rolland et al. 2023). The primary aim of these interventions is to reduce age-related morbidity or mortality and to maintain function. Ideally, such interventions begin long before pathology leads to a notable decline. However, clinical trials spanning the many years or decades needed to observe the effect on human aging would be very difficult and expensive. Aging biomarkers have been proposed as a means for researchers to assess the impact of specific interventions within a feasible time frame for clinical studies (Moqri et al. 2023; Aging Biomarker Consortium et al. 2023). Such biomarkers are trained to quantify biological age or pace of aging as a proxy for longer-term outcomes. However, research into how these biomarkers respond to interventions, and the significance of observed biomarker changes, remains in its infancy.

Epigenetic clocks are aging biomarkers based on DNA methylation at cytosine-guanine dinucleotides (CpGs). These clocks have gained significant popularity over the past decade due to their prognostic power and the ease and speed of measurement, requiring a simple blood draw (Horvath & Raj 2018; Drew 2022). The first generation of epigenetic clocks, such as the Hannum (Hannum et al. 2013), Horvath multi-tissue (Horvath MT)(Horvath 2013), and Horvath skin-and-blood (Horvath SB) clocks (Horvath & Raj 2018), utilized penalized regression models (e.g., elastic net) to predict chronological age from DNA methylation patterns. Newer generations of clocks use similar techniques but are trained to predict mortality and morbidity risk; these include PhenoAge (Levine et al. 2018) and GrimAge (Lu et al. 2019). Another recent clock model, DunedinPoAm38, was trained on longitudinal biomarkers to predict an individual’s pace of aging (Belsky et al. 2020). Many of these epigenetic-based measurements of biological age have been shown to be prognostic, correlating at least partially with outcomes such as mortality (Simpson & Chandra 2021).

Beyond predictive capabilities, epigenetic clocks may be used in a clinical trial setting to rapidly calculate an individual’s biological age before and after an aging intervention and assess the impact of treatment. Epigenetic clocks have already been applied in this fashion for several interventions, such as diet, exercise, and supplements (Sae-Lee et al. 2018; Gensous et al. 2020; Fitzgerald et al. 2021; Yumi Noronha et al. 2022). However, there are several potential issues with using epigenetic clocks as metrics in longitudinal trials.

The first potential problem with measuring the impact of an intervention on epigenetic age is longitudinal reliability. We previously demonstrated that re-testing the same individual, either by testing the same sample multiple times, or by conducting testing at multiple follow-up time points, can lead to fluctuations by several years owing to technical noise and other confounders (Higgins-Chen et al. 2022). This concern led to the development of PC clocks (Higgins-Chen et al. 2022), re-trained versions of the canonical clocks mentioned above that use principal component analysis to identify age-related patterns across a larger number of CpGs and reduce the effect of noise from individual CpGs. These PC clock variants reduce longitudinal variability for a single individual, increasing our ability to reliably detect the impact of an intervention on biological age while reducing false positives. Similarly, DunedinPACE is a modified version of the DunedinPoAm38 pace-of-aging predictor that increases reliability and longitudinal performance by only utilizing reliable CpGs as input (Belsky et al. 2022).

While the development of reliable clock models may help with measuring biological age longitudinally, the existence of multiple unique clock models leads to additional practical issues that need to be addressed before the clocks can be used in a clinical setting. With an abundance of clock models, which one should a researcher select for their particular study? How can we be sure which clock is the most relevant? And if multiple clocks are calculated for a study, how do we interpret the situation where different clocks disagree? This is an ongoing problem in the literature - several studies that use epigenetic clocks to measure an aging intervention report results from a single clock model in their analysis (Sae-Lee et al. 2018; Gensous et al. 2020; Fitzgerald et al. 2021; Yumi Noronha et al. 2022) but it is unclear if the chosen clock is most appropriate. Under these conditions, the field is at risk for publication bias - opting for clocks that return significant results and ignoring the non-significant results. A direct comparison of the responsiveness of the various clocks to aging interventions is warranted to help correct this issue.

We hypothesize that some significant epigenetic clock changes are not replicable using any other clock model because they are false positives due to noise. Meanwhile, significant clock changes that are replicable across multiple clocks are *bona fide* changes in epigenetic age. To investigate this hypothesis, we calculate 6 well-established epigenetic clocks along with their high-reliability counterparts, for 10 publicly available longitudinal DNA methylation datasets. Eight of these datasets contain methylation data before and after a proposed aging intervention. We focus on diet, exercise, and lifestyle studies to increase comparability between studies. To act as positive controls, we analyze two datasets that capture an event likely to increase the biological age of subjects (i.e., cancer treatment or intensive surgery), reasoning it should be easier to accelerate aging than decelerate it. Our study shows that some clock changes are likely the result of technical noise and false positives, and provides guidelines for selecting combinations of clocks and multiple testing correction to increase the likelihood that an epigenetic clock change reflects a valid aging intervention effect.

## Results

### False Positives: Multiple intervention studies show sporadic changes in chronological-age clocks

We calculated the change in subject biological age residual (Δ resid, see methods) using 12 epigenetic clocks for 6 publicly available datasets, before and after an intervention (Figure 1, Table 1 and 2). The interventions we examined included acupuncture (GSE184202), daily supplements (GSE63499, GSE74538), high intensity exercise (GSE171140), or a combination of diet and lifestyle changes (GSE149747, E-MTAB-8956). Datasets GSE149474 and GSE74538 were associated with studies that previously reported changes in the Horvath MT clock, but did not examine any other clocks in their analysis (Fitzgerald et al. 2021; Sae-Lee et al. 2018). The timeframes for the studies chosen here varied in range from a few hours to 2 years. Details on the datasets and studies selected for this analysis can be found in the Methods section. We performed a Student’s paired t-test on epigenetic age residual (Methods) before and after the intervention for each of the 12 epigenetic clocks (Figure 1). In datasets which had control cohorts, we also calculated unpaired t-tests between the subject and control groups age residual but found no significant changes (Supplemental Figure 1). Our initial analyses do not employ multiple testing correction, given that we are probing the possibilities of false positive results in these datasets, and are not rejecting a null hypothesis based on the significance of a single t-test. Additionally, as most studies do not employ multiple clocks, it is not well-established which method of multiple hypothesis correction is appropriate. Later, we investigate the impact of multiple hypothesis testing on our results (Table 3).

**Figure 1:**
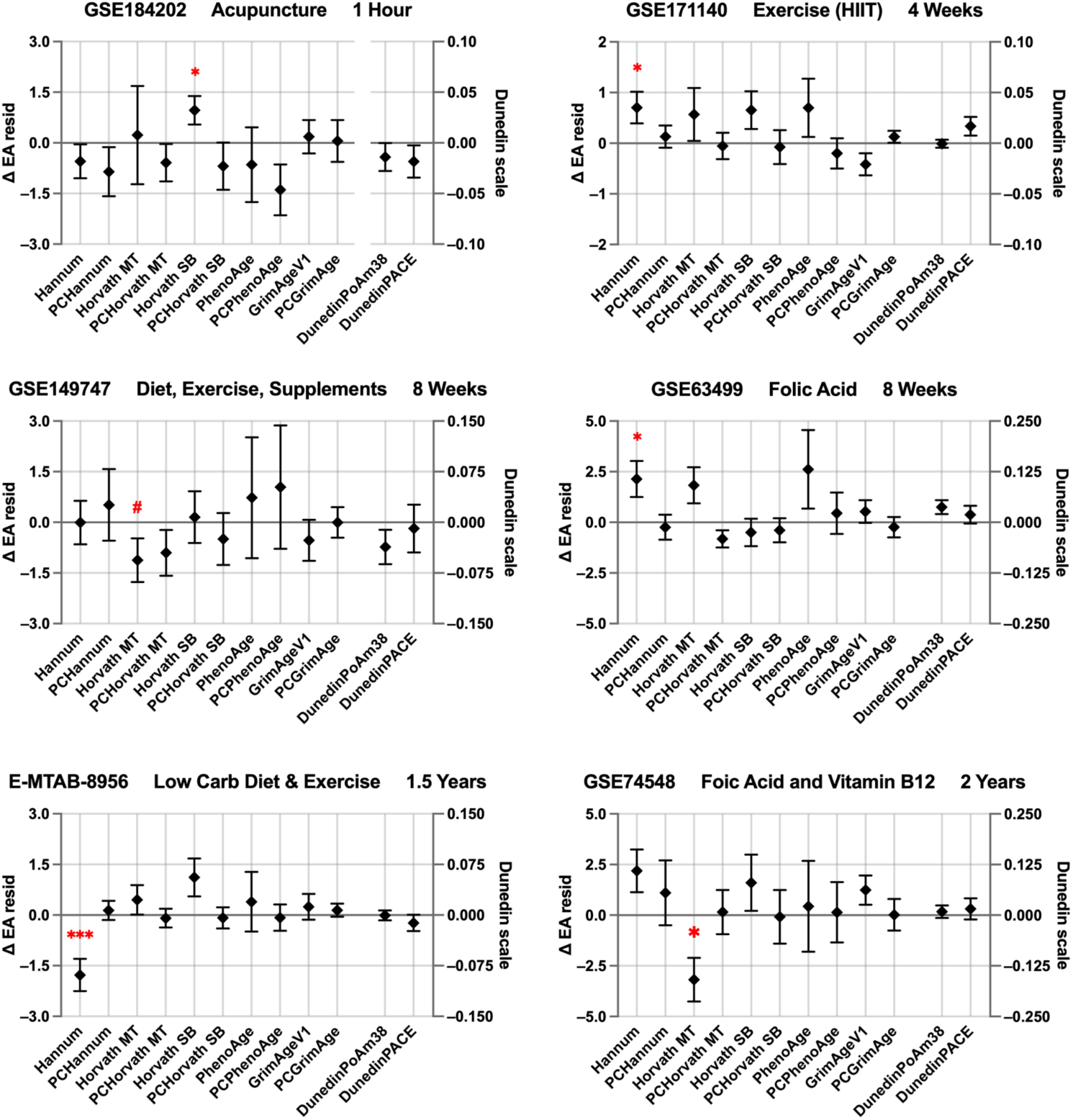
Twelve epigenetic clock models measuring six aging intervention datasets with sporadic significance. Black diamonds represent the mean change in epigenetic age residual (Δ resid) for all subjects in the cohort. Upper and lower black error bars indicate standard error of the mean. Secondary y-axis (right) resolution is increased by 20x for DunedinPoAm38 and DunedinPACE. Red asterisk indicates a significant result for that clock for that dataset, as calculated by a paired t-test (p value < 0.05). Red hash for GSE149747 indicates a p-value of 0.066.

**Table 1:** Summary of the publicly available human DNA methylation datasets which had multiple timesteps per subject (longitudinal) and which captured an intervention or event during the trial.

| Accession # | Reference | Intervention / Event Type | Duration | N |
| --- | --- | --- | --- | --- |
| GSE184202 | (Petitpierre et al. 2022) | Acupuncture | Hours | 11 |
| GSE171140 | (Voisin et al. 2020) | High Intensity Interval Training Exercise | 4 weeks | 36 |
| GSE149747 | (Fitzgerald et al. 2021) | Diet, Supplements, Exercise, and Lifestyle | 8 weeks | 19 |
| GSE63499 | (Shade et al. 2017) | Folic Acid Supplements | 8 weeks | 12 |
| E-MTAB-8956 | (Yaskolka Meir et al. 2016; Gepner et al. 2018) | Low Carb Diet and Exercise | 1.5 years | 30 |
| GSE74548 | (Sae-Lee et al. 2018) | Folic Acid and Vitamin B12 Supplements | 2 years | 14 |
| GSE142536 | (Sadahiro et al. 2020) | Intensive Surgery | 1 day | 30 |
| GSE140038 | (Sehl et al. 2020) | Radiotherapy with or without Chemotherapy | Months | 72 |
| E-MTAB-12527 | (Yaskolka Meir et al. 2021) | Mediterranean Diet with Red Meat | 2 Years | 81 |
| E-MTAB-12527 | (Yaskolka Meir et al. 2021) | Mediterranean Diet with No Red Meat | 2 Years | 87 |

In 5 of the 6 longitudinal intervention datasets, there was a single clock which found a significant change in epigenetic age residual (either decreasing or increasing). This includes GSE74538, which previously reported a significant change in Horvath MT (Sae-Lee et al. 2018). The remaining dataset, GSE149747, showed a trend towards reduction in one clock, Horvath MT (p=0.066), consistent with previously published results (Fitzgerald et al. 2021). In all cases, the lone clock that reported a significant result was a first-generation clock, which had been trained to measure chronological age. In no cases did the PC version of these clocks corroborate the significant result. In 3 of the 6 datasets, the significant change in biological age is positive, suggesting that these interventions actually increase biological age – something which seems counterintuitive, given the known health benefits of these interventions. Their increase is more consistent with our hypothesis that these sporadically significant findings are a result of a type-1 error. Even if the sporadic result is a *bona fide* change in a particular clock, the fact that no other clock shows any similar effect raises the question about the biological significance of the result.

### Positive Control: Age-accelerating events are captured by multiple reliable clocks

If it is possible to capture an intervention that decreases biological age using epigenetic clocks, then it stands to reason that the reverse should be true: events that are known to increase mortality, and increase risk of death, should result in biological age acceleration. Indeed, a prior study showed that stressful events (surgery, pregnancy, severe COVID-19) lead to strong but reversible increases in epigenetic age according to multiple clocks (Poganik et al. 2023). We reasoned that we could treat these age-accelerating events as positive controls. By observing their effects on epigenetic clocks, we can gain insight into what would constitute a trustworthy pattern of epigenetic clock changes in response to aging interventions. With this hypothesis in mind, we repeated our 12-clock analysis on two longitudinal datasets that captured events with a known association with mortality (Figure 2, Table 1). We examined epigenetic clocks before and after intensive surgery (GSE142536, previously analyzed by Poganik et al. 2023) as well as before and after radiation and chemotherapy (GSE140038, not previously analyzed).

**Figure 2:**
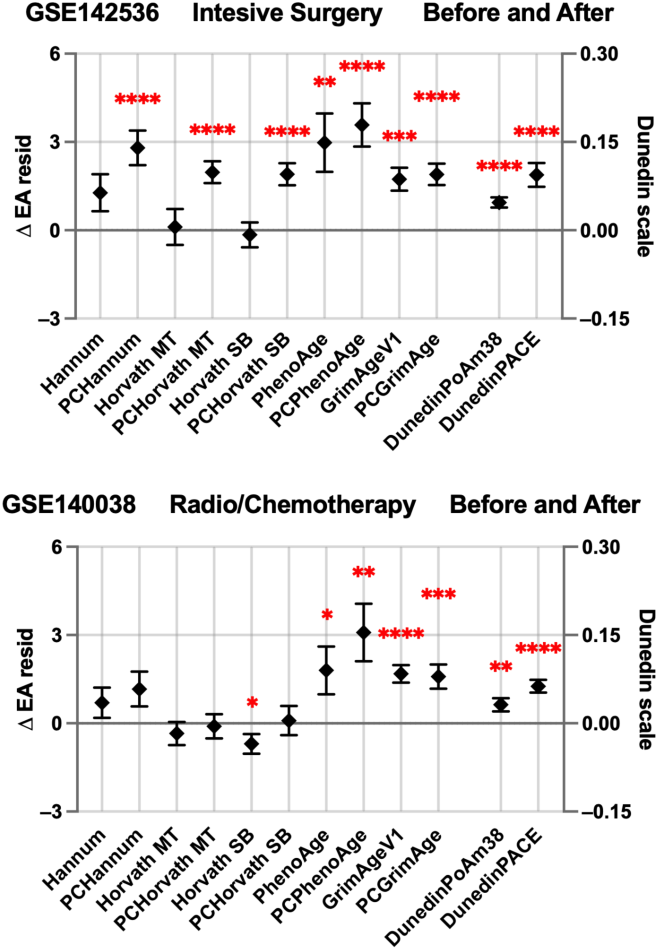
Positive control datasets, capturing age accelerating events, measured with 12 epigenetic clock models. Black diamonds indicate the average change in epigenetic age residual (Δ resid) amongst subjects. Error bars indicate standard error of the mean. Red asterisks indicate a significant change, as measured by a paired t-test.

In both datasets, we found significant increases in at least 6 of the 12 clocks that we tested (Figure 2). All the mortality-based clocks (PhenoAge, GrimAge), their PC analogs, and the rate of aging clocks (DunedinPoAm38, DunedinPACE) agreed on a significant increase in biological age residual after the event. In the dataset comparing biological age before and after intensive surgery, all PC versions of the clocks captured a significant increase in biological age, while the standard versions of the chronological based clocks did not see a significant change.

Despite the intensity of the events that the subjects underwent, the chronological based clocks (Hannum, Horvath MT, and Horvath SB) fail to report a significant increase in biological age. In fact, Horvath2 indicates a significant decrease in biological age after radiotherapy and chemotherapy. Taken together, this reinforces our finding that the chronological trained clocks are poor proxies for measuring aging interventions. Instead, the high-reliability clocks, as well as clocks predicting mortality or pace of aging, are better suited for detecting intervention effects.

### True Positives: Validated lifestyle interventions modify reliable and Gen 2 clock

The insights from the previous analysis on positive control datasets suggests interventions that impact aging should be present in multiple clocks, including the mortality, rate of aging, and PC variant clocks. We identified a single longitudinal aging intervention study that showed this type of epigenetic clock response. We repeated our 12-clock analysis for a 2-year diet trial (E-MTAB-12527) with two arms. One cohort ate a standard Mediterranean-style diet (MED) and another cohort ate a Mediterranean diet with more red meat restrictions and enriched with green plants and polyphenols (green) (Figure 3). We examined changes in epigenetic age after each dietary intervention.

**Figure 3:**
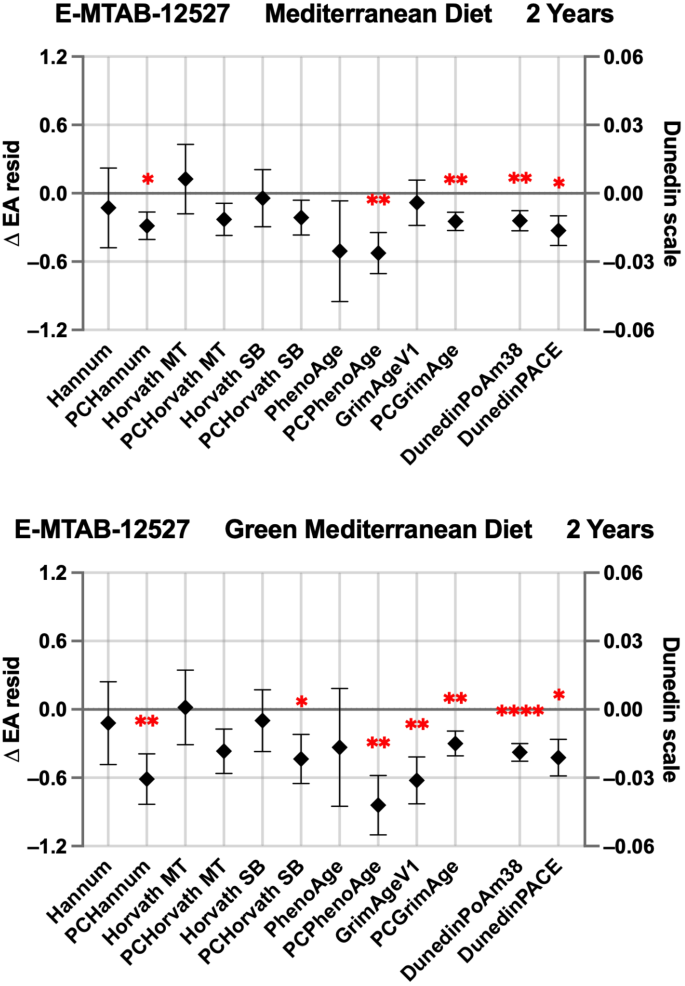
Two examples of two year-long Mediterranean diets significantly reducing multiple different epigenetic clock models, including reliable (PC), mortality-trained, and pace-of-aging clocks. Black diamonds indicate mean change in epigenetic age residual (Δ resid) for a particular clock model. Error bars represent standard error of the mean. Red asterisks indicate a significant change for that clock model, as measured by a paired t-test.

We found a significant decrease of biological age in 5 and 7 of the 12 epigenetic clocks in the MED and green diet, respectively (Figure 3). These significant decreases present in one of the three clock groups: PC clocks, mortality-based clocks, and rate-of-aging clocks. None of the first-generation chronological-based non-PC clocks reported a decrease in biological age. These results mirror the positive control results in Figure 2. Therefore, we suggest the epigenetic clock results in this study are indeed capturing some positive health benefits related to aging and are sharply distinct from the false positive results in Figure 1.

### True Positives, but not False Positives, pass multiple testing correction

We hypothesized that the sporadic significant results found in the datasets in Figure 1 are the result of multiple hypothesis testing. That is, repeating 12 t-tests on different metrics of the same dataset compounds the risk of a false positive result. If this is the case, then correcting for multiple comparisons should remove those type-1 errors. Likewise, if the effect we are seeing in the positive controls (Figure 2) and true positive results (Figure 3) are genuine responses to the interventions, and not statistical noise, then they should remain even after the testing correction.

We applied the Bonferroni and Benjamini-Hochberg methods in each study separately to evaluate the impact of multiple comparisons. The Bonferroni method adjusts the family-wide error rate, by dividing the p value threshold for significance by the number of statistical tests (in our case, 12), and is therefore the most stringent method. The Benjamini-Hochberg method ranks the p values and sets a dynamic critical threshold, where the smallest p value receives the strictest test (essentially a Bonferroni) and the largest p value receives the most lenient threshold (the standard 0.05 false discovery rate).

We find that 4 of the 5 initial datasets with sporadic significance in the non-PC chronological based clocks lose their significance after multiple hypothesis correction with either Bonferroni or Benjamini-Hochberg (Table 3, life-style intervention GSE171140; acupuncture GSE184202; folic acid supplements GSE63499; and folic acid and vitamin B12 supplement GSE74548). Compare this to the positive control datasets (GSE140038, GSE142536) --most of the clocks (8 out of 9 for GSE142536 and 4 out of 7 for GSE140038) remain significant even with the more stringent multiple hypothesis correction (Bonferroni). Similarly, our hypothesized “true positive” interventions (E-MTAB-12527) remain statistically significant in all but one clock after Benjamini-Hochberg. Of note, the clocks that did not pass Benjamini-Hochberg correction in these positive control or true positive interventions tended to be lower-reliability clocks (original PhenoAge or DunedinPoAm38) or chronological age clocks (PCHannum, PCHorvathSB, HorvathSB). In contrast, high-reliability mortality clocks like GrimAge, PCGrimAge, PCPhenoAge, or DunedinPACE were much more likely to pass multiple testing correction. This brief analysis validates our hypothesis that a single significant clock after an intervention is likely a false positive, whereas multiple highly significant clocks that stand up to multiple hypothesis correction suggest a genuine intervention-based impact on the biological mechanisms of aging.

## Discussion

Epigenetic clocks represent a promising biomarker candidate for assessing the impact of an aging intervention. However, not all clocks are designed the same, and the ability to respond to an aging intervention is not necessarily conserved across all epigenetic clock models.

In the datasets we analyzed, clocks trained on chronological age, i.e. first-generation clocks (Hannum, Horvath MT, and Horvath SB), failed to concur with any other clock models on the impact of an aging intervention. They almost always responded alone, and they often failed to respond when their PC variant or multiple other clock models did detect a significant change. This observation suggests that first-generation clocks, while accurate at predicting chronological age, are inaccurate for detecting biological age changes and therefore they should not be used to assess the impact of an intervention. This result is not surprising when you consider how the clock models were trained – to predict chronological age. This training process prioritizes methylation sites that are more dependent on time, and less dependent on additional confounders, such as lifestyle or a particular diet or supplement. Furthermore, their lower reliability means that first-generation clock changes are more likely the result of technical noise rather than *bona fide* changes in biological age.

Moreover, recent findings suggest that epigenetic age may fluctuate as much as 2 years during the course of a single day (Koncevičius et al. 2024). This inherent rhythmicity may be introducing false positives when relying on single clock tests, as the daily variation can be misinterpreted as an intervention effect. This further underscores the need for more reliable models that are less vulnerable to time-of-day, but also other potential confounders such as fasting status, acute stress, menstrual cycles, time-of-year, etc.

Conversely, the clocks trained on mortality, or the pace of aging clocks DunedinPoAm38 and DunedinPACE, only indicated a significant change in biological age in concert with other clocks. This agreement between the clock models regarding the impact of a particular intervention or aging event reaffirms our confidence in their results. In contrast to the first-generation clocks, these clocks are trained to predict aging outcomes and it is reassuring that they respond, in unison, to aging interventions and events. It is likely that the methylation sites that these clocks use to predict age have more relevance to health and aging hallmarks than those sites used in the first-generation clocks.

Our finding that the non-PC first generation clocks respond sporadically and unreliably to a range of aging interventions has implications for past, ongoing, and future clinical aging interventions trials that use one or more epigenetic clocks. One significant clock is not enough to indicate a reliable decrease in biological age, especially if the PC variant of that clock fails to show a significant trend. There are several studies, already published, that recognize this concern *a priori* and utilize multiple epigenetic clocks in their analysis. The impact of calorie restrictions (Waziry et al. 2023) and umbilical cord plasma transfusions (Clement et al. 2022) on biological age have both been investigated using multiple epigenetic clocks, providing more nuance in the interpretation of their intervention’s impact on biological age. However, this is not the norm, and even recent intervention studies that use and report multiple clocks will interpret one positive result from a chronological clock, while ignoring the mortality-trained, or pace-of-aging, clocks that show no significant change (da Silva Rodrigues et al. 2024; Patterson et al. n.d.).

This selective reporting of positive results raises concerns about potential publication bias. Researchers may unintentionally favor clocks that show significant changes, even if other, more reliable clocks do not. This bias highlights the need for a more holistic approach where a battery of clocks is tested simultaneously to avoid overinterpreting the result of a single, potentially unreliable clock. If an intervention is decreasing biological age, the change should register with more than one epigenetic clock, ideally a later generation reliable clock model such as the PC clocks, PhenoAge, GrimAge, and DunedinPACE.

The use of multiple, diverse epigenetic clock models to assess the impact of an aging intervention or event is critical, as it significantly reduces the chance of interpreting a stray result as a genuine reduction in biological age. Here, we present one possible battery of clocks to apply to any longitudinal intervention study, that contains a diverse variety of models. This multi-clock approach provides researchers with a more nuanced understanding of the impact of an intervention, as each clock was trained slightly differently, and each therefore measures a slightly different definition of biological age. This method will help to bolster confidence in the use of epigenetic clock models for measuring aging interventions, and will drive future clinical trial development aimed at extending human healthspan.

## Methods

### Data Acquisition and Preprocessing

Where available, DNA methylation data were downloaded as beta values from public repositories, specifically the Gene Expression Omnibus (GEO) database (NIH) or the European Molecular Biology Laboratory’s European Bioinformatics Institute (EMBL-EBI). For datasets where methylation beta values were not directly available (E-MTAB-8956, E-MTAB-12527), we retrieved the raw fluorescence intensity files (idat files). These raw files were subsequently processed and converted into methylation beta values using the minfi package in R (version 1.48.0), following Normalization of Oligonucleotide Arrays by Background Subtraction (NOOB) and Quantile Normalization. DNA Methylation datasets used either Illlumina 450k or Illumina 850k array platforms.

Phenotypic data was not directly modified, rather, six additional curated columns were appended to the phenotypic data frame. The six columns were adapted from the source data, and included sample ID, subject ID, sex, age, group (control vs subject), and time (in days). This step was done for all datasets, to ensure standardized and replicable data handling in downstream analysis.

In cases of missing methylation beta values, mean imputation was performed within the subject cohort. Missing beta values (NAs) were replaced with the average beta value for all individuals in the cohort, ensuring a complete dataset for downstream analysis.

### Datasets

For datasets containing multiple timepoints, only two timepoints were selected for the analysis: the pre-intervention baseline sample and a post-intervention follow-up sample. For these datasets, the time point selected for the follow-up sample was always the first follow-up time point. This approach was applied to maintain consistency and reduce complexity in longitudinal comparisons. Dataset GSE74548 was subset to include only female participants with the MTHFR 677CC genotype, aligning with the significant findings reported by (Sae-Lee et al. 2018). Intensive surgery, for the case of dataset GSE142536, includes elective colorectal surgery, elective hip replacement surgery, and emergency hip surgery following fracture (Sadahiro et al. 2020).

### Epigenetic Clock Calculation

We calculated scores for 12 well-established epigenetic clocks, as summarized in Table 2. These clocks include both first-generation clocks trained to predict chronological age (e.g., Hannum, Horvath MT, Horvath SB) and newer generation clocks trained to predict mortality risk or rate of aging (e.g., GrimAge, PhenoAge, DunedinPACE). We also calculated the PC version of these clocks, where available. Epigenetic clock scores for the two Horvath clocks, Hannum, PhenoAge, and DunedinPoAm38 were computed from the methylation beta values using the MethylCIPHER R package (version 0.20, https://github.com/HigginsChenLab/methylCIPHER). PC clock scores were calculated using the PC clocks package (https://github.com/HigginsChenLab/PC-Clocks). GrimAgeV1 was calculated with a custom R function adapted from the biolearn python package (https://bio-learn.github.io/). DunedinPACE was calculated using the DunedinPACE R package (version 0.99.0, https://github.com/danbelsky/DunedinPACE).

**Table 2:** Summary of the epigenetic clock models, trained to predict either chronological age, mortality risk, or pace of aging. The PC clocks represent a group of clocks, including PCHannum, PCHorvath MT, PCHorvath SB, PCPhenoAge, and PCGrimAge.

| Clock | Trained to predict... | # of CpGs | Reference |
| --- | --- | --- | --- |
| Hannum | Chronological Age | 71 | (Hannum et al. 2013) |
| Horvath multi-tissue (MT) | Chronological Age | 353 | (Horvath 2013) |
| Horvath skin and blood (SB) | Chronological Age | 391 | (Horvath et al. 2018) |
| PhenoAge | Mortality Risk | 513 | (Levine et al. 2018) |
| GrimAge | Mortality Risk | 1,030 | (Lu et al. 2019) |
| DunedinPoAm38 | Pace of aging | 47 | (Belsky et al. 2020) |
| DunedinPACE | Pace of aging | 173 | (Belsky et al. 2022) |
| PC clocks | Chronological Age/Mortality Risk | 78,464 | (Higgins-Chen et al. 2022) |

### Statistical analysis

The primary outcome measure, age residual, was calculated for each subject by regressing predicted epigenetic age (DNAmAge) on chronological age by using the following linear model:

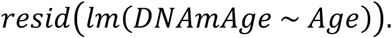

Models were built in R using the stats package (version 4.3.2). Paired t-tests were conducted to compare age residuals before and after the intervention, paired by subject ID, also in R using the stats package. The change in age residual (Δ resid) for one subject across the intervention was computed as follows:

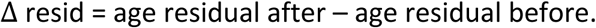

In Figures 1-3, DunedinPoAm38 and DunedinPACE are plotted against a separate y-axis (on the right) which was scaled to 1/20 of the left y-axis. This was done to better visualize the smaller absolute outputs from those clocks.

### Multiple Testing Correction

We performed Bonferroni and Benjamini-Hochberg corrections to account for multiple hypothesis testing (Table 3). Calculations were performed in R using a custom-built function. Scripts for multiple hypothesis correction can be found at GitHub.

**Table 3:**
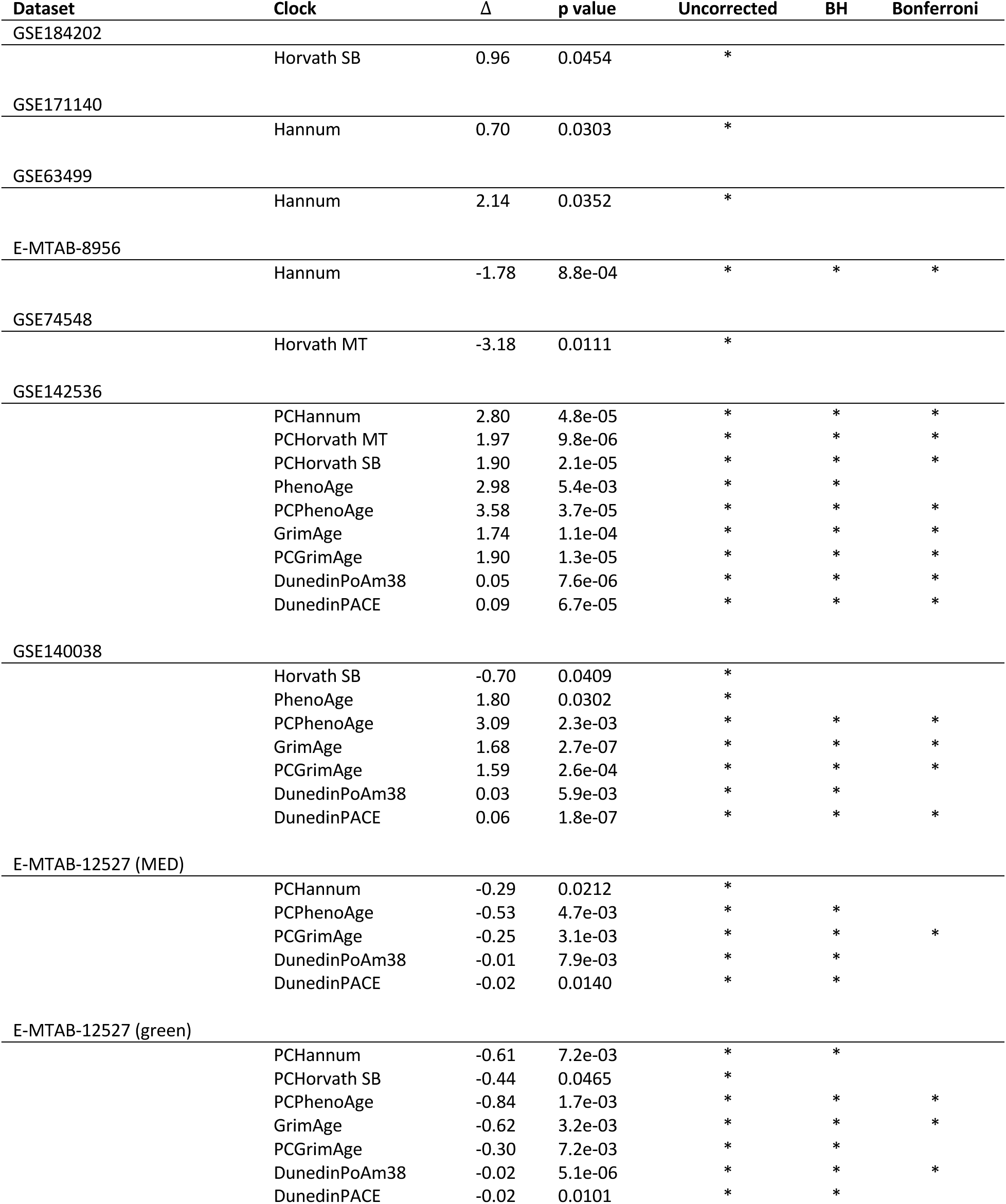
Results from multiple hypothesis correction for 12 statistical tests. Δ indicates the mean change in epigenetic age residual from before to after intervention/event. Uncorrected column has no multiple hypothesis correction. BH: Benjamini-Hochberg. An asterisk represents a significant result for that statistical test with the particular hypothesis correction method as defined by the column.

| Dataset | Clock | $\Delta$ | p value | Uncorrected | BH | Bonferroni |
| --- | --- | --- | --- | --- | --- | --- |
| GSE184202 | Horvath SB | 0.96 | 0.0454 | * |  |  |
| GSE171140 | Hannum | 0.70 | 0.0303 | * |  |  |
| GSE63499 | Hannum | 2.14 | 0.0352 | * |  |  |
| E-MTAB-8956 | Hannum | -1.78 | 8.8e-04 | * | * | * |
| GSE74548 | Horvath MT | -3.18 | 0.0111 | * |  |  |
| GSE142536 | PCHannum | 2.80 | 4.8e-05 | * | * | * |
|  | PCHorvath MT | 1.97 | 9.8e-06 | * | * | * |
|  | PCHorvath SB | 1.90 | 2.1e-05 | * | * | * |
|  | PhenoAge | 2.98 | 5.4e-03 | * | * |  |
|  | PCPhenoAge | 3.58 | 3.7e-05 | * | * | * |
|  | GrimAge | 1.74 | 1.1e-04 | * | * | * |
|  | PCGrimAge | 1.90 | 1.3e-05 | * | * | * |
|  | DunedinPoAm38 | 0.05 | 7.6e-06 | * | * | * |
|  | DunedinPACE | 0.09 | 6.7e-05 | * | * | * |
| GSE140038 | Horvath SB | -0.70 | 0.0409 | * |  |  |
|  | PhenoAge | 1.80 | 0.0302 | * |  |  |
|  | PCPhenoAge | 3.09 | 2.3e-03 | * | * | * |
|  | GrimAge | 1.68 | 2.7e-07 | * | * | * |
|  | PCGrimAge | 1.59 | 2.6e-04 | * | * | * |
|  | DunedinPoAm38 | 0.03 | 5.9e-03 | * | * |  |
|  | DunedinPACE | 0.06 | 1.8e-07 | * | * | * |
| E-MTAB-12527 (MED) | PCHannum | -0.29 | 0.0212 | * |  |  |
|  | PCPhenoAge | -0.53 | 4.7e-03 | * | * |  |
|  | PCGrimAge | -0.25 | 3.1e-03 | * | * | * |
|  | DunedinPoAm38 | -0.01 | 7.9e-03 | * | * |  |
|  | DunedinPACE | -0.02 | 0.0140 | * | * |  |
| E-MTAB-12527 (green) | PCHannum | -0.61 | 7.2e-03 | * | * |  |
|  | PCHorvath SB | -0.44 | 0.0465 | * |  |  |
|  | PCPhenoAge | -0.84 | 1.7e-03 | * | * | * |
|  | GrimAge | -0.62 | 3.2e-03 | * | * | * |
|  | PCGrimAge | -0.30 | 7.2e-03 | * | * |  |
|  | DunedinPoAm38 | -0.02 | 5.1e-06 | * | * | * |
|  | DunedinPACE | -0.02 | 0.0101 | * | * |  |

## Acknowledgements

The authors would like to thank the clinicians and researchers who made their study data publicly available, as without their data this project would not have been possible. The authors would also like to thank everyone in the Higgins-Chen lab for their invaluable feedback and insights pertaining to this project.

## Author contributions

A.H.C., D.S.B., R.S. conceived the project and study design. D.S.B. developed the analysis pipeline and identified relevant datasets, and R.S., J.F.A., J.K., and J.G. assisted in dataset and code preparation. A.H.C. and D.S.B. wrote the manuscript. All authors helped edit and prepare the manuscript for submission.

## Data availability statement

All methylation array data in this study comes from publicly available sources. All effects sizes will be posted upon publication. Code to calculate all clocks is accessible at https://github.com/HigginsChenLab/methylCIPHER.

---

**Supplemental Figure 1:**
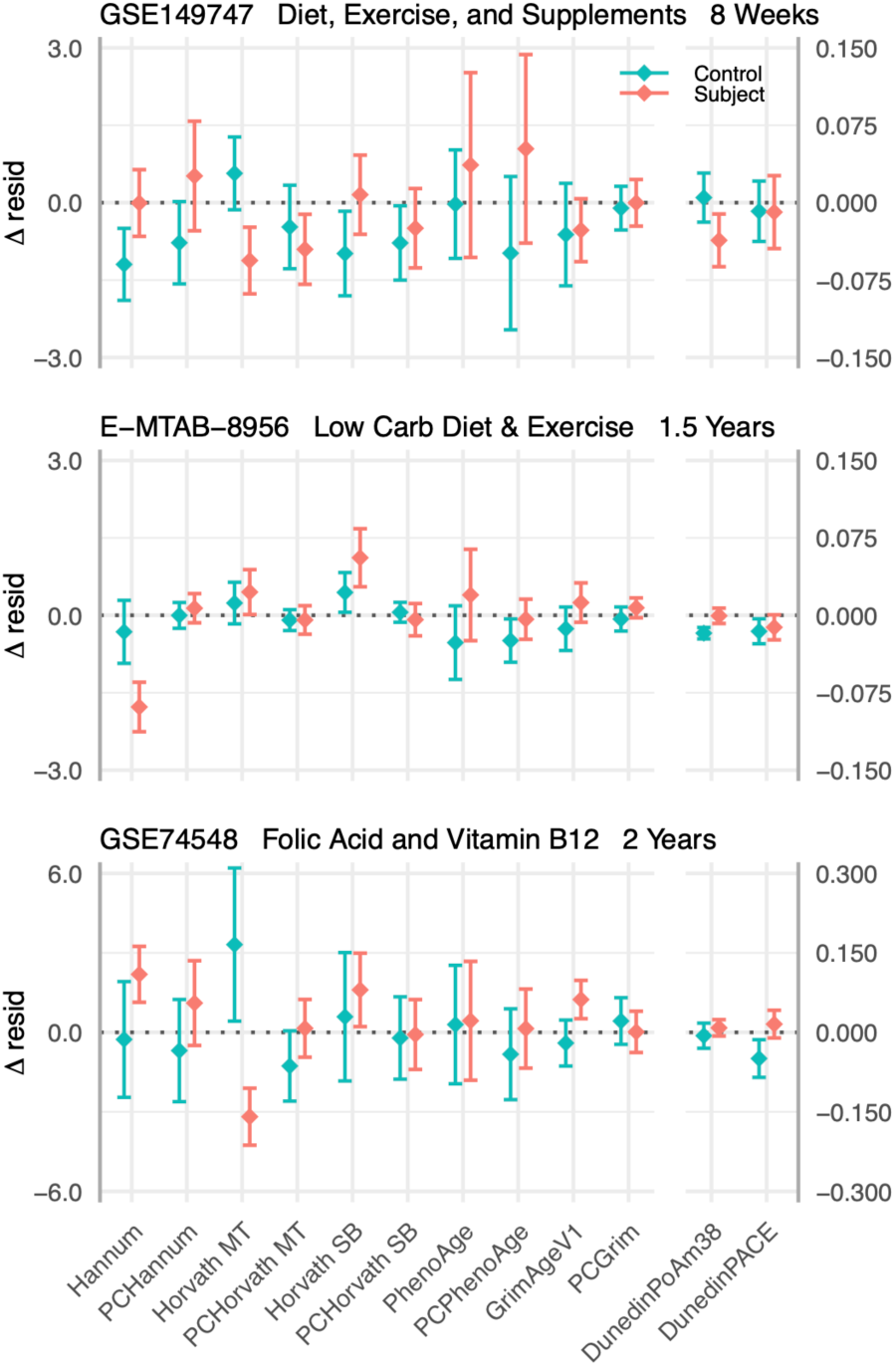
Comparison of changes in epigenetic age residuals (Δ resid) across various interventions. Diamonds represent the average change in age residual for control (cyan) and subject (pink) groups. Error bars represent standard error of the mean.

## References

Aging Biomarker Consortium, Bao H, Cao J, Chen M, Chen M, Chen W, Chen X, Chen Y, Chen Y, Chen Y, Chen Z, Chhetri JK, Ding Y, Feng J, Guo J, Guo M, He C, Jia Y, Jiang H, Jing Y, Li D, Li J, Li J, Liang Q, Liang R, Liu F, Liu X, Liu Z, Luo OJ, Lv J, Ma J, Mao K, Nie J, Qiao X, Sun X, Tang X, Wang J, Wang Q, Wang S, Wang X, Wang Y, Wang Y, Wu R, Xia K, Xiao F-H, Xu L, Xu Y, Yan H, Yang L, Yang R, Yang Y, Ying Y, Zhang L, Zhang W, Zhang W, Zhang X, Zhang Z, Zhou M, Zhou R, Zhu Q, Zhu Z, Cao F, Cao Z, Chan P, Chen C, Chen G, Chen H-Z, Chen J, Ci W, Ding B-S, Ding Q, Gao F, Han J-DJ, Huang K, Ju Z, Kong Q-P, Li J, Li J, Li X, Liu B, Liu F, Liu L, Liu Q, Liu Q, Liu X, Liu Y, Luo X, Ma S, Ma X, Mao Z, Nie J, Peng Y, Qu J, Ren J, Ren R, Song M, Songyang Z, Sun YE, Sun Y, Tian M, Wang S, Wang S, Wang X, Wang X, Wang Y-J, Wang Y, Wong CCL, Xiang AP, Xiao Y, Xie Z, Xu D, Ye J, Yue R, Zhang C, Zhang H, Zhang L, Zhang W, Zhang Y, Zhang Y-W, Zhang Z, Zhao T, Zhao Y, Zhu D, Zou W, Pei G & Liu G-H (2023) Biomarkers of aging. Sci. China Life Sci. 66, 893–1066.

Belsky DW, Caspi A, Arseneault L, Baccarelli A, Corcoran DL, Gao X, Hannon E, Harrington HL, Rasmussen LJ, Houts R, Huffman K, Kraus WE, Kwon D, Mill J, Pieper CF, Prinz JA, Poulton R, Schwartz J, Sugden K, Vokonas P, Williams BS & Moffitt TE (2020) Quantification of the pace of biological aging in humans through a blood test, the DunedinPoAm DNA methylation algorithm. Elife 9. Available at: 10.7554/eLife.54870.

Belsky DW, Caspi A, Corcoran DL, Sugden K, Poulton R, Arseneault L, Baccarelli A, Chamarti K, Gao X, Hannon E, Harrington HL, Houts R, Kothari M, Kwon D, Mill J, Schwartz J, Vokonas P, Wang C, Williams BS & Moffitt TE (2022) DunedinPACE, a DNA methylation biomarker of the pace of aging. Elife 11. Available at: 10.7554/eLife.73420.

Clement J, Yan Q, Agrawal M, Coronado RE, Sturges JA, Horvath M, Lu AT, Brooke RT & Horvath S (2022) Umbilical cord plasma concentrate has beneficial effects on DNA methylation GrimAge and human clinical biomarkers. Aging Cell 21, e13696.

Drew L (2022) Turning back time with epigenetic clocks. Nature 601, S20–S22.

Fitzgerald KN, Hodges R, Hanes D, Stack E, Cheishvili D, Szyf M, Henkel J, Twedt MW, Giannopoulou D, Herdell J, Logan S & Bradley R (2021) Potential reversal of epigenetic age using a diet and lifestyle intervention: a pilot randomized clinical trial. Aging (Albany NY) 13, 9419–9432.

Gensous N, Garagnani P, Santoro A, Giuliani C, Ostan R, Fabbri C, Milazzo M, Gentilini D, di Blasio AM, Pietruszka B, Madej D, Bialecka-Debek A, Brzozowska A, Franceschi C & Bacalini MG (2020) One-year Mediterranean diet promotes epigenetic rejuvenation with country-and sex-specific effects: a pilot study from the NU-AGE project. GeroScience 42, 687–701.

Hannum G, Guinney J, Zhao L, Zhang L, Hughes G, Sadda S, Klotzle B, Bibikova M, Fan J-B, Gao Y, Deconde R, Chen M, Rajapakse I, Friend S, Ideker T & Zhang K (2013) Genome-wide methylation profiles reveal quantitative views of human aging rates. Mol. Cell 49, 359–367.

Higgins-Chen AT, Thrush KL, Wang Y, Minteer CJ, Kuo P-L, Wang M, Niimi P, Sturm G, Lin J, Moore AZ, Bandinelli S, Vinkers CH, Vermetten E, Rutten BPF, Geuze E, Okhuijsen-Pfeifer C, van der Horst MZ, Schreiter S, Gutwinski S, Luykx JJ, Picard M, Ferrucci L, Crimmins EM, Boks MP, Hägg S, Hu-Seliger TT & Levine ME (2022) A computational solution for bolstering reliability of epigenetic clocks: Implications for clinical trials and longitudinal tracking. *Nat*. Aging 2, 644–661.

Horvath S (2013) DNA methylation age of human tissues and cell types. Genome Biol. 14, R115.

Horvath S & Raj K (2018) DNA methylation-based biomarkers and the epigenetic clock theory of ageing. Nat. Rev. Genet. 19, 371–384.

Koncevičius K, Nair A, Šveikauskaitė A, Šeštokaitė A, Kazlauskaitė A, Dulskas A & Petronis A (2024) Epigenetic age oscillates during the day. *Aging Cell*, e14170.

Levine ME, Lu AT, Quach A, Chen BH, Assimes TL, Bandinelli S, Hou L, Baccarelli AA, Stewart JD, Li Y, Whitsel EA, Wilson JG, Reiner AP, Aviv A, Lohman K, Liu Y, Ferrucci L & Horvath S (2018) An epigenetic biomarker of aging for lifespan and healthspan. Aging (Albany NY) 10, 573–591.

López-Otín C, Blasco MA, Partridge L, Serrano M & Kroemer G (2023) Hallmarks of aging: An expanding universe. Cell 186, 243–278.

Lu AT, Quach A, Wilson JG, Reiner AP, Aviv A, Raj K, Hou L, Baccarelli AA, Li Y, Stewart JD, Whitsel EA, Assimes TL, Ferrucci L & Horvath S (2019) DNA methylation GrimAge strongly predicts lifespan and healthspan. Aging (Albany NY) 11, 303–327.

Moqri M, Herzog C, Poganik JR, Biomarkers of Aging Consortium, Justice J, Belsky DW, Higgins-Chen A, Moskalev A, Fuellen G, Cohen AA, Bautmans I, Widschwendter M, Ding J, Fleming A, Mannick J, Han J-DJ, Zhavoronkov A, Barzilai N, Kaeberlein M, Cummings S, Kennedy BK, Ferrucci L, Horvath S, Verdin E, Maier AB, Snyder MP, Sebastiano V & Gladyshev VN (2023) Biomarkers of aging for the identification and evaluation of longevity interventions. Cell 186, 3758–3775.

Patterson W, Rossner RJ, Garuda R, Davis M & Terry GC Plasmid delivery of follistatin gene therapy safely improves body composition and lowers extrinsic epigenetic age in sex-and age-diverse adult human subjects. Available at: https://minicircle.io/wp-content/uploads/2024/04/fstpreprint.pdf.

Poganik JR, Zhang B, Baht GS, Tyshkovskiy A, Deik A, Kerepesi C, Yim SH, Lu AT, Haghani A, Gong T, Hedman AM, Andolf E, Pershagen G, Almqvist C, Clish CB, Horvath S, White JP & Gladyshev VN (2023) Biological age is increased by stress and restored upon recovery. Cell Metab. 35, 807– 820.e5.

Rolland Y, Sierra F, Ferrucci L, Barzilai N, De Cabo R, Mannick J, Oliva A, Evans W, Angioni D, De Souto Barreto P, Raffin J, Vellas B, Kirkland JL & G.C.T-TF group (2023) Challenges in developing Geroscience trials. Nat. Commun. 14, 5038.

Sadahiro R, Knight B, James F, Hannon E, Charity J, Daniels IR, Burrage J, Knox O, Crawford B, Smart NJ & Mill J (2020) Major surgery induces acute changes in measured DNA methylation associated with immune response pathways. Sci. Rep. 10, 5743.

Sae-Lee C, Corsi S, Barrow TM, Kuhnle GGC, Bollati V, Mathers JC & Byun H-M (2018) Dietary intervention modifies DNA methylation age assessed by the epigenetic clock. Mol. Nutr. Food Res. 62, e1800092.

da Silva Rodrigues G, Noma IHY, Noronha NY, Watanabe LM, da Silva Sobrinho AC, de Lima JGR, Sae-Lee C, Benjamim CJR, Nonino CB & Bueno CR Júnior (2024) Eight weeks of physical training decreases 2 years of DNA methylation age of sedentary women. Res. Q. Exerc. Sport 95, 405–415.

Simpson DJ & Chandra T (2021) Epigenetic age prediction. Aging Cell 20, e13452.

Waziry R, Ryan CP, Corcoran DL, Huffman KM, Kobor MS, Kothari M, Graf GH, Kraus VB, Kraus WE, Lin DTS, Pieper CF, Ramaker ME, Bhapkar M, Das SK, Ferrucci L, Hastings WJ, Kebbe M, Parker DC, Racette SB, Shalev I, Schilling B & Belsky DW (2023) Effect of long-term caloric restriction on DNA methylation measures of biological aging in healthy adults from the CALERIE trial. *Nat*. Aging 3, 248–257.

Yumi Noronha N, da Silva Rodrigues G, Harumi Yonehara Noma I, Fernanda Cunha Brandao C, Pereira Rodrigues K, Colello Bruno A, Sae-Lee C, Moriguchi Watanabe L, Augusta de Souza Pinhel M, Mello Schineider I, Luciano de Almeida M, Barbosa Júnior F, Araújo Morais D, Tavares de Sousa Júnior W, Plösch T, Roberto Bueno Junior C & Barbosa Nonino C (2022) 14-weeks combined exercise epigenetically modulated 118 genes of menopausal women with prediabetes. Front. Endocrinol. (Lausanne) 13, 895489.

